## Supplemental Figures for "Crosshair, semi-automated targeting for electron microscopy with a motorised ultramicrotome"

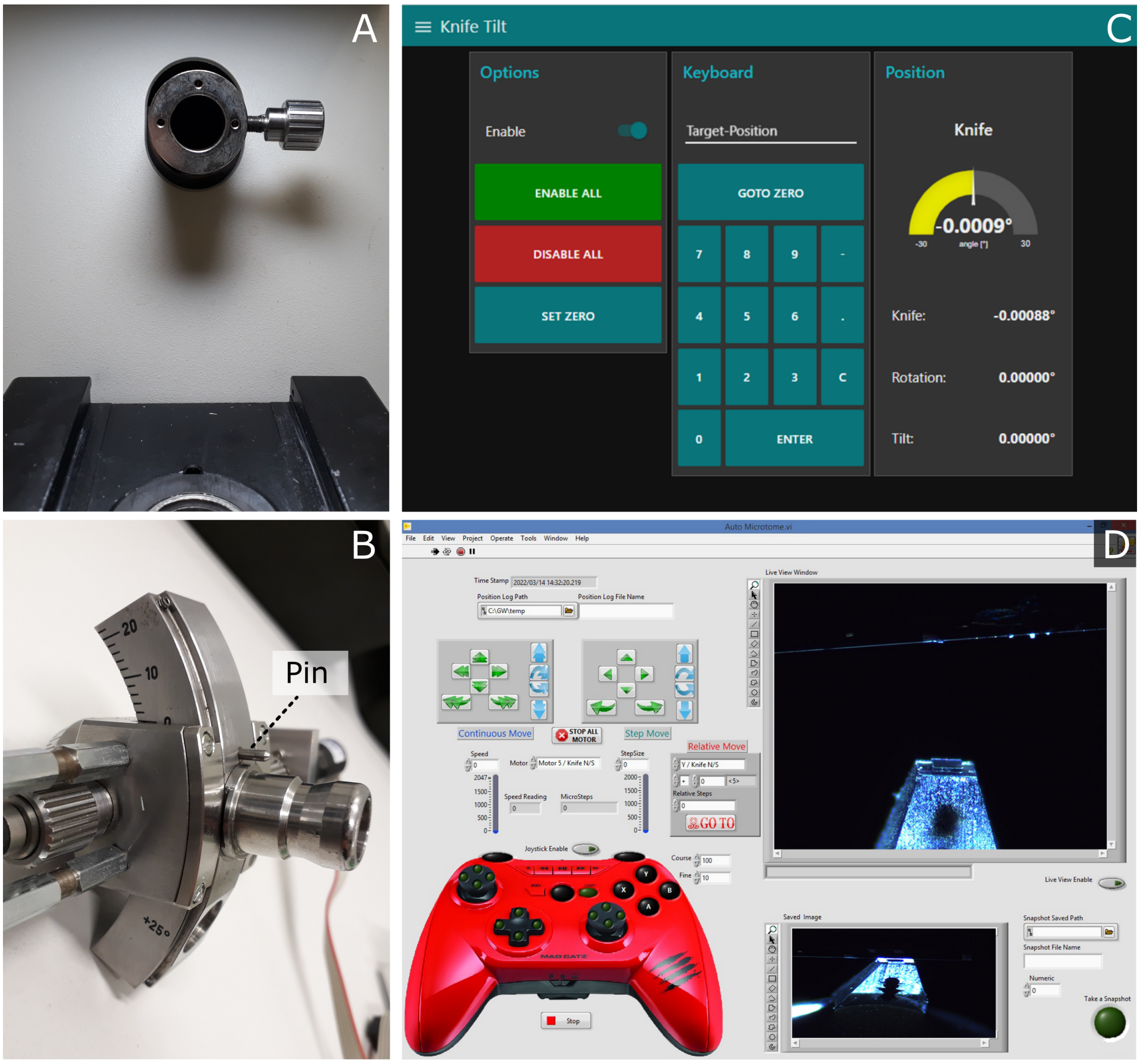


**Figure S1:** Motorised ultramicrotome. **A** - Leica ultramicrotome arm with sample holder removed. This shows the additional hole that was drilled (here there are 3 holes, but the top one is the only one required) **B** - zoom of the back of the Leica sample holder, showing the small metal pin that was added to fit into the hole in *A*. **C** - Knife tilt page of the user interface for the motorised Leica system. This is shown on a touchscreen to the right of the ultramicrotome, and allows motors to be enabled/disabled and set to specific angles. **D** - user interface of the motorised RMC system.

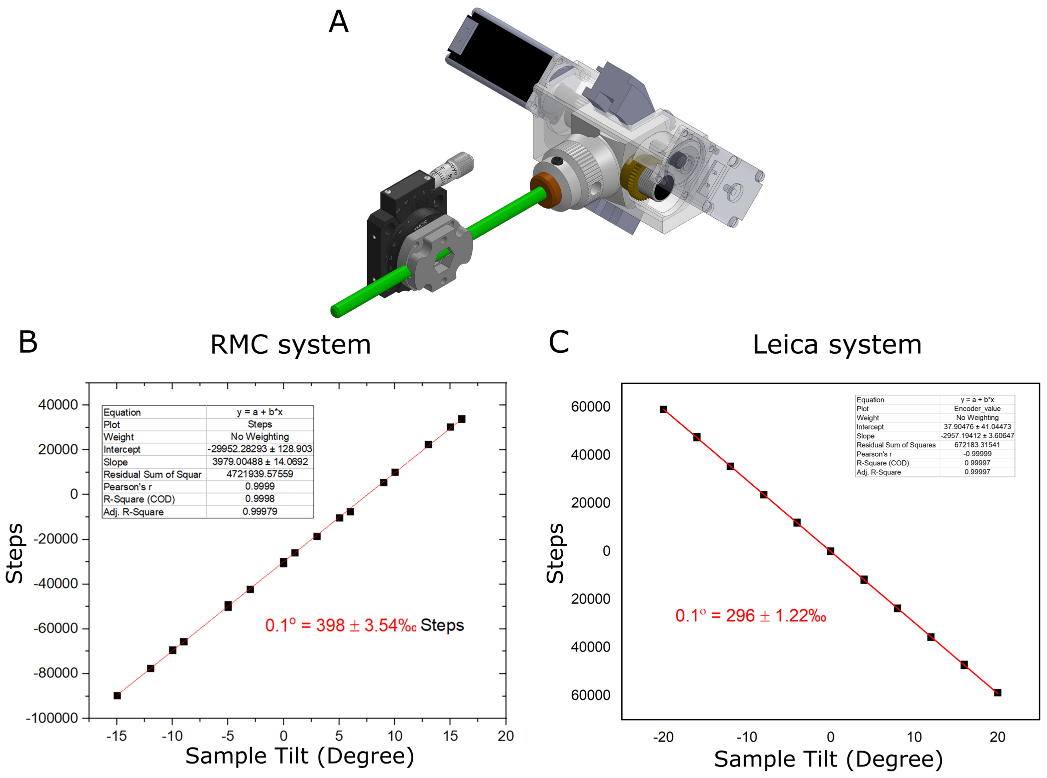


**Figure S2:** Calibration of sample tilt axis. **A** - Diagram of calibration setup, with RMC system. **B** - graph of sample tilt vs motor steps for RMC system. A linear fit is shown in red. **C** - Same as *B* for Leica system.

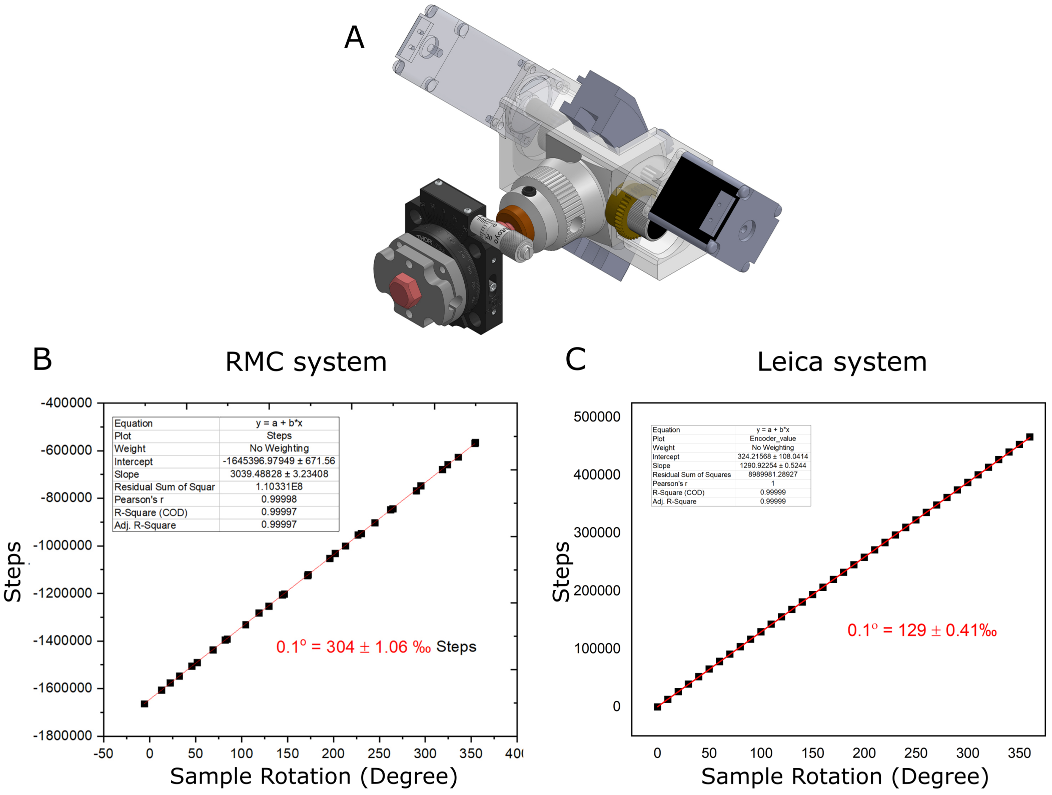


**Figure S3:** Calibration of sample rotation axis. **A** - Diagram of calibration setup, with RMC system. **B** - graph of sample rotation vs motor steps for RMC system. A linear fit is shown in red. **C** - Same as *B* for Leica system.

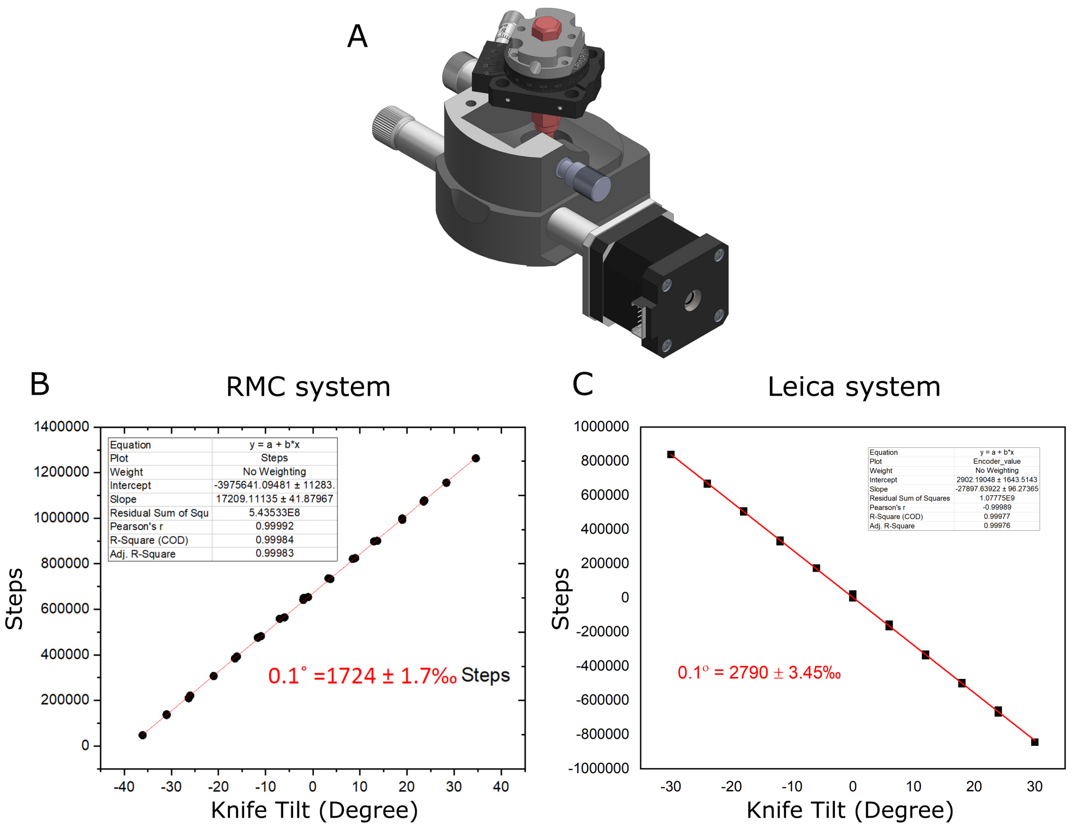


**Figure S4:** Calibration of knife tilt axis. **A** - Diagram of calibration setup, with RMC system. **B** - graph of knife tilt vs motor steps for RMC system. A linear fit is shown in red. **C** - Same as *B* for Leica system.

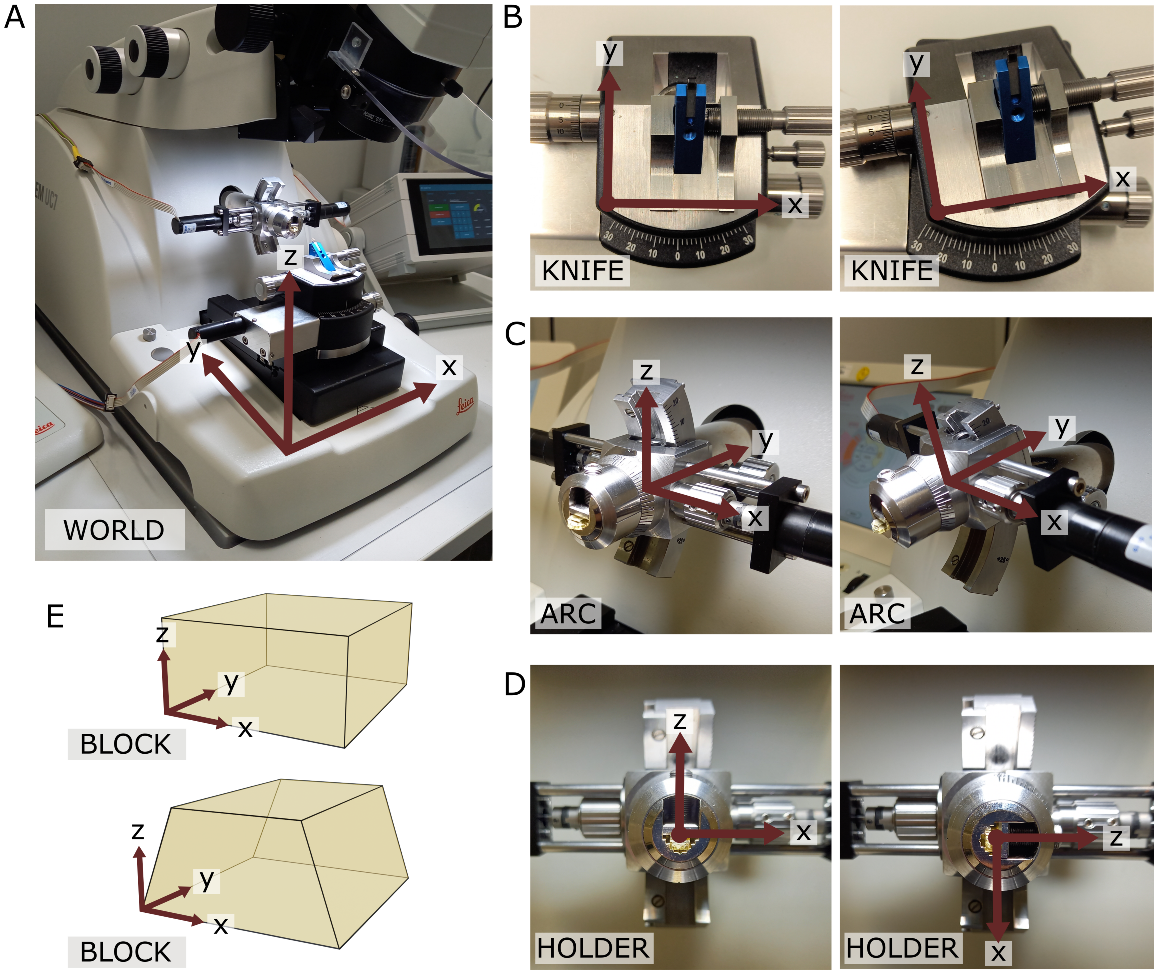


**Figure S5:** Coordinate frames. **A** - world coordinate frame. X is parallel to EW, y is parallel to NS, z is vertical. **B** - Knife coordinate frame at 0 degrees (left) and 10 degrees (right). Z points out of the page, and is parallel to the world coordinate frame z. Note that at 0 degrees the knife frame has the same orientation as the world frame. **C** - Arc coordinate frame at 0 degrees (left) and 14 degrees (right). X is parallel to the world coordinate frame x. Note that at 0 degrees the arc frame has the same orientation as the world frame. **D** - Holder coordinate frame at two positions separated by a 90 degree rotation. Note that here the 0 position depends on the particular targeting run, as it is defined as the rotation when the knife and block face are aligned. The y axis points into the page and is parallel to the arc coordinate frame y axis. **E** - Block coordinate frame for a rectangular block face (top) and a trapezium block face (bottom). X is parallel to the bottom edge of the block face.

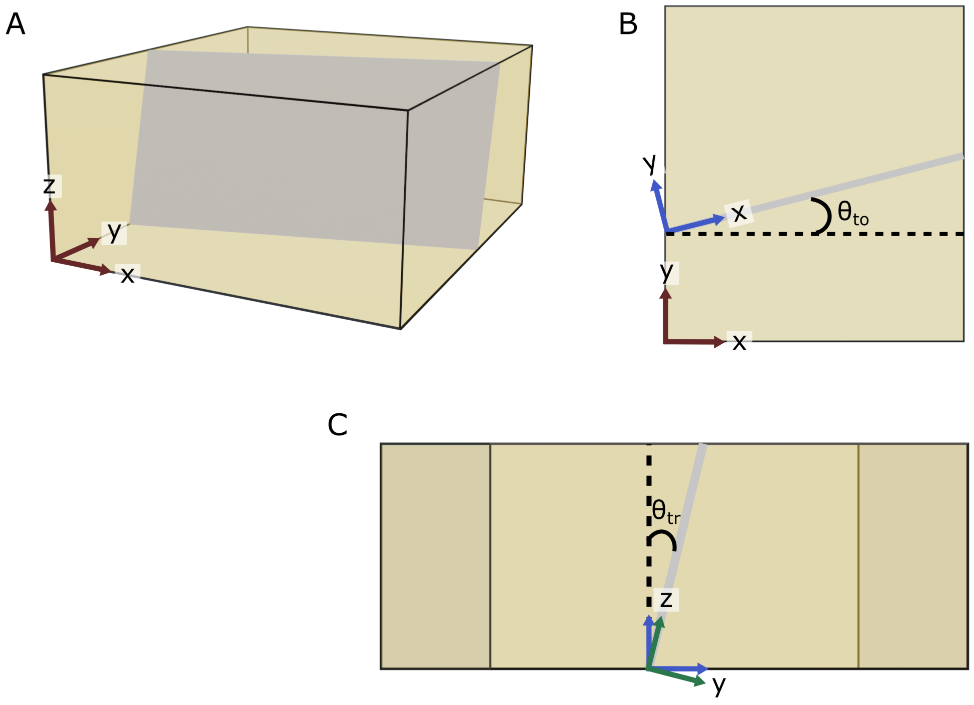


**Figure S6:** Block and target coordinate frames. **A** - diagram of a rectangular block, with the block coordinate frame shown in red. The gray plane inside the block is the target plane. **B** - Considering the xy plane only, the target plane intersects along the shown gray line. $\theta_{to}$ is then defined as the angle between the block frame x axis and the blue x axis shown i.e. the rotation angle about the block frame z. **C** - Now considering the zy plane of the blue axes from *B* (i.e. looking down the line of intersection), $\theta_{tr}$ is defined as the angle between the blue and green z axes i.e. the rotation angle about the blue frame x. The green axes are the final target coordinate frame.

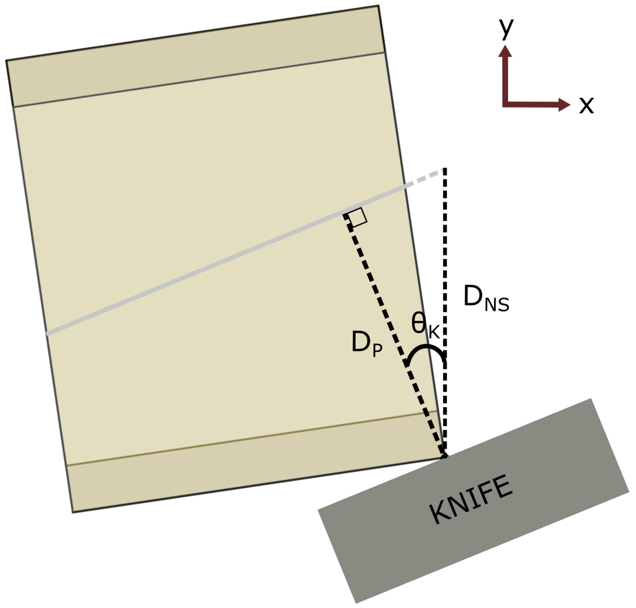


**Figure S7:** Distance calculation. Here, the same block and target plane as in figure S6 are shown in an example solution orientation. In red is the world coordinate frame, showing we are looking down the world z axis. The knife is aligned to the vertical target plane, but the ultramicrotome arm (and therefore the block) advances towards the knife in the world y axis (NS) between cuts. Therefore, the required cutting distance is not the same as the perpendicular distance between the knife and target plane. $D_P$ is the perpendicular distance between the target plane and the furthest surface point, $D_{NS}$ is the corresponding NS distance (i.e the required cutting depth), and $\theta_K$ is the knife angle.

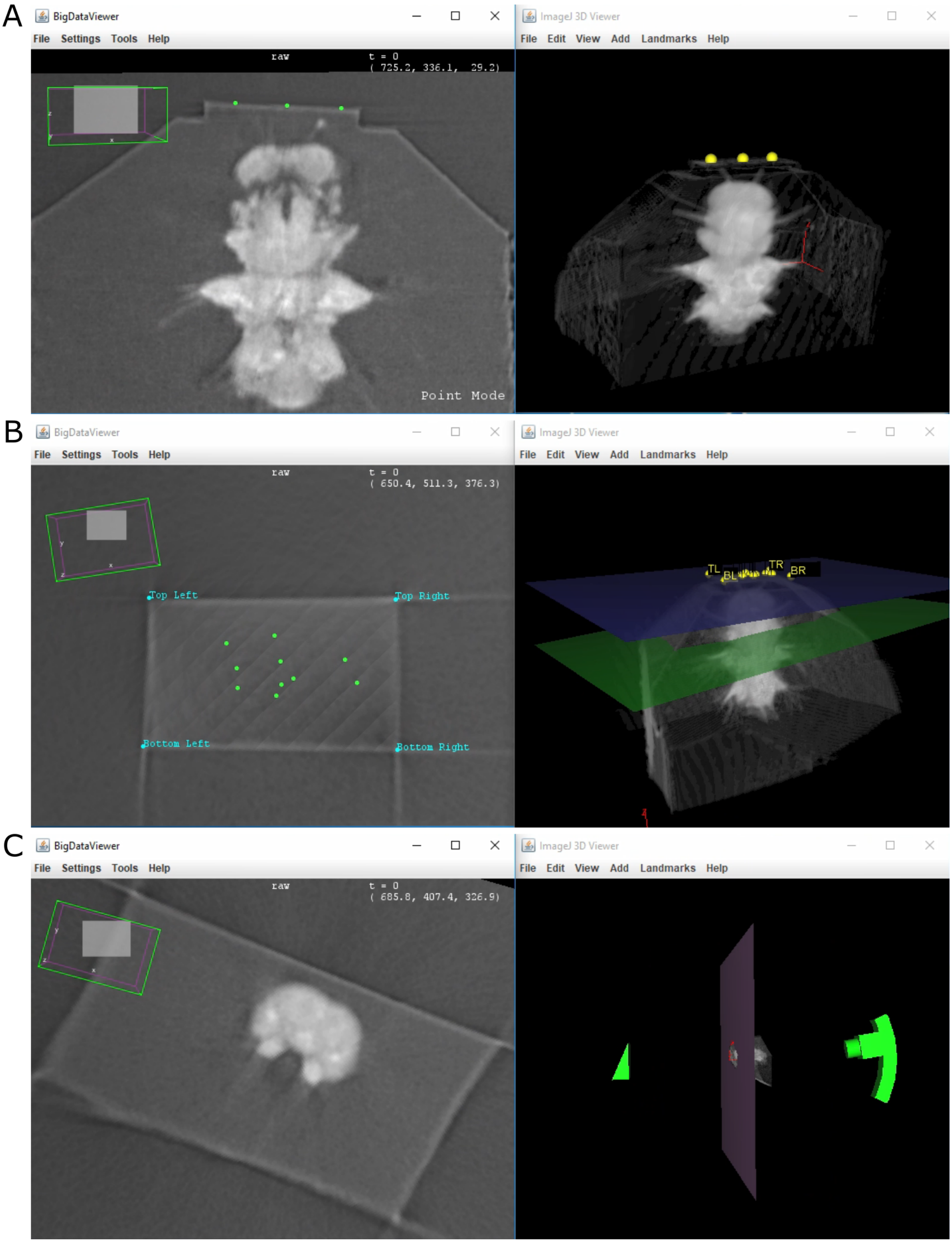


**Figure S8:** Crosshair. Left column - 2D slice. Right column - 3D volume rendering. **A** - 3 points placed on the block surface to fit a plane. **B** - block surface shown in 2D and 3D (blue plane) with labelled corners and orientation. The green points in the 2D view are the points used to fit the plane originally. **C** - example of 'cutting mode'. The 2D slice shows the predicted surface at a certain cut depth, indicated in 3D by the large pink plane.

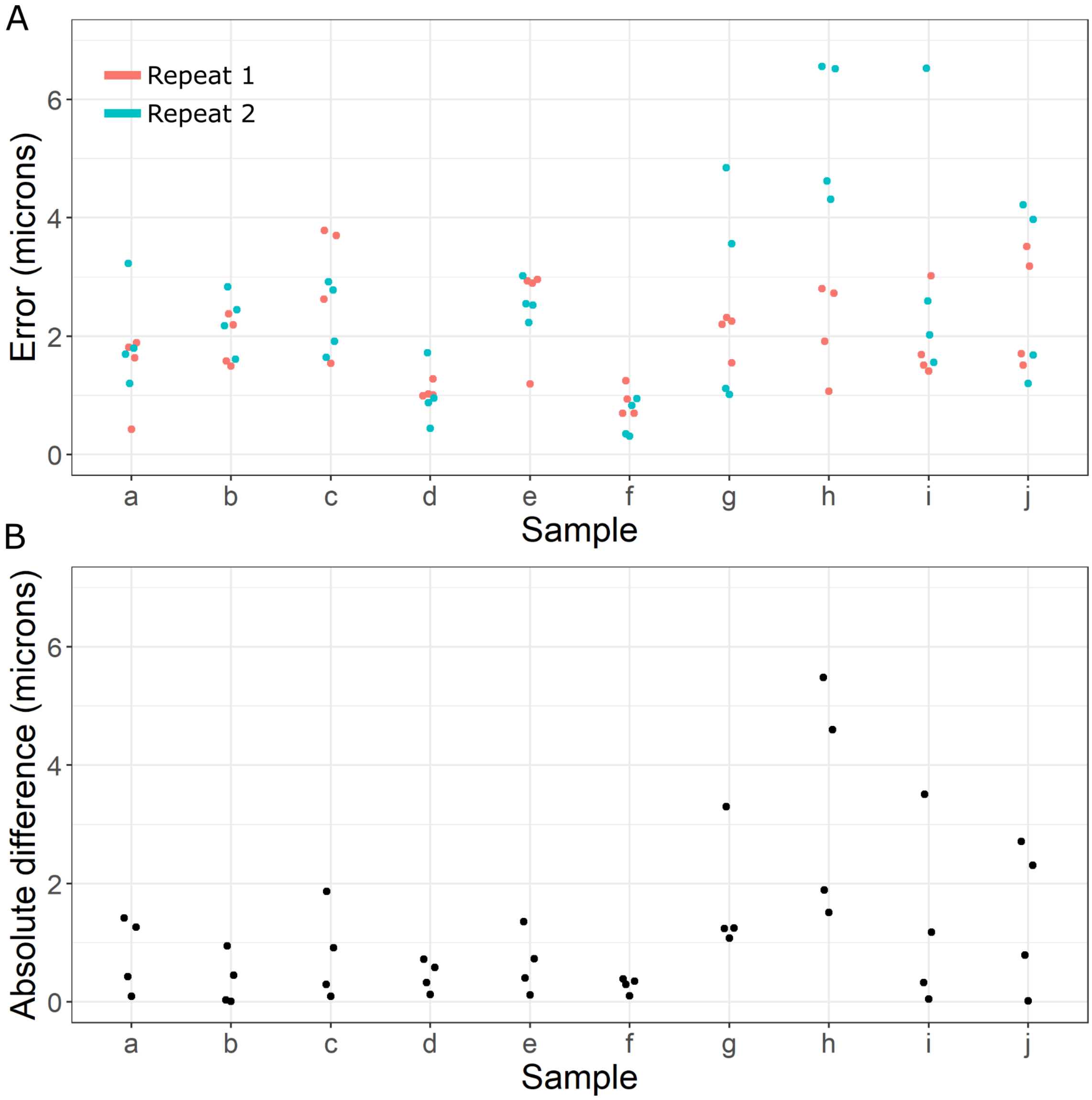


**Figure S9:** Registration accuracy. **A** - Beeswarm plot of distance error (microns) measured between manually placed corresponding points on pairs of registered X-ray images for samples a-j. Two repeats of manually placing points in the same locations, on the same images are shown. **B** - absolute difference (microns) between the distance error measured in *A* between repeat 1 and repeat 2 for each point.

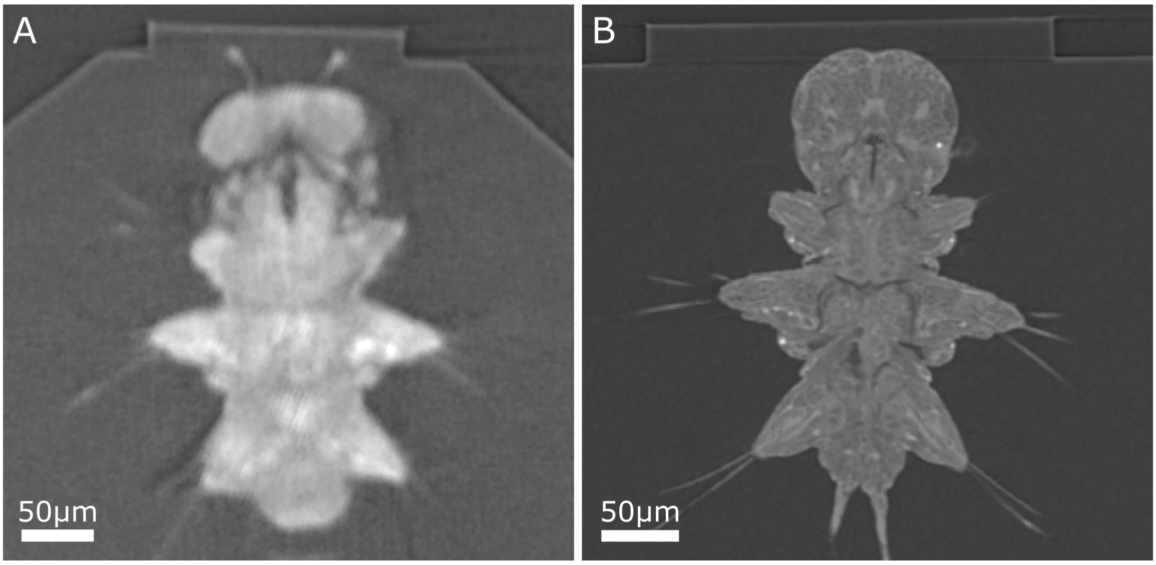


**Figure S10:** Comparison of example *Platynereis* X-ray cross-sections, both acquired with 1 micron isotropic voxel size. **A** - Sample d X-ray, acquired with Bruker SkyScan 1272 (system used for samples a-e). **B** - Sample f X-ray, acquired with Zeiss Versa 510 (system used for samples f-j). Note the sharper block edges and the finer anatomical details visible in *B*.

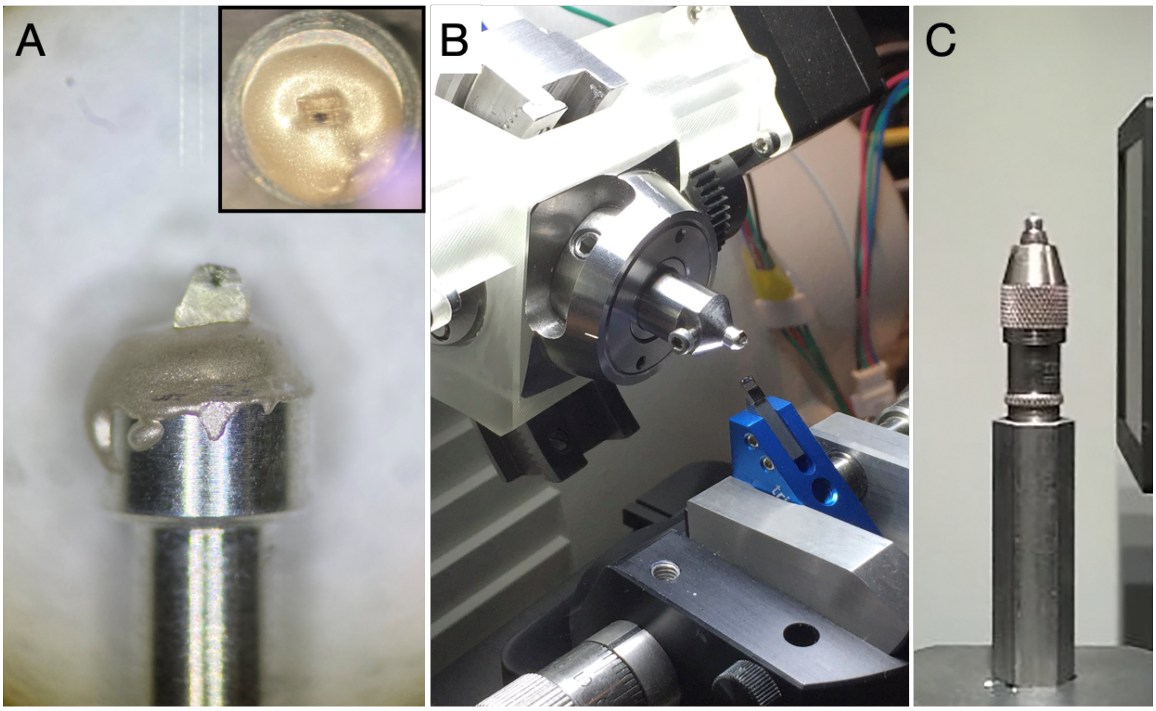


**Figure S11:** Sample mounting for trimming, X-ray imaging and targeting for samples f-j. **A** - Resin-embedded sample mounted onto an aluminium pin using conductive epoxy glue. **B** - An aluminium pin with sample inserted into a Gatan Rivet Holder which is directly inserted into an ultramicrotome chuck holder for trimming and targeting at the ultramicrotome. **C** - An aluminium pin with sample mounted to a sample holder of Versa 510 X-ray microscope for X-ray imaging.
